# Whole-brain, all-optical interrogation of neuronal dynamics underlying gut interoception in zebrafish

**DOI:** 10.1101/2025.03.26.645305

**Authors:** Weiyu Chen, Ben James, Virginia M. S. Ruetten, Sambashiva Banala, Ziqiang Wei, Greg Fleishman, Mikail Rubinov, Mark C. Fishman, Florian Engert, Luke D. Lavis, James E. Fitzgerald, Misha B. Ahrens

## Abstract

Internal signals from the body and external signals from the environment are processed by brain-wide circuits to guide behavior. However, the complete brain-wide circuit activity underlying interoception—the perception of bodily signals—and its interactions with sensorimotor circuits remain unclear due to technical barriers to accessing whole-brain activity at the cellular level during organ physiology perturbations. We developed an all-optical system for whole-brain neuronal imaging in behaving larval zebrafish during optical uncaging of gut-targeted nutrients and visuo-motor stimulation. Widespread neural activity throughout the brain encoded nutrient delivery, unfolding on multiple timescales across many specific peripheral and central regions. Evoked activity depended on delivery location and occurred with amino acids and D-glucose, but not L-glucose. Many gut-sensitive neurons also responded to swimming and visual stimuli, with brainstem areas primarily integrating gut and motor signals and midbrain regions integrating gut and visual signals. This platform links body-brain communication studies to brain-wide neural computation in awake, behaving vertebrates.

## Introduction

The importance of interoceptive signals from body organs in information processing in the brain, and the critical role of the brain in controlling the physiological state of the body, have long been appreciated^1–12^. Through body-brain interactions, hunger causes profound changes in mood and decision making^13–15^, and sickness causes drastic changes in behavior beyond muscle weakness^16^. These effects on cognition and behavior imply widespread changes to neuronal activity and underscore the need for a whole-brain, cellular-resolution approach to studying interoception. However, despite rapid progress in our understanding of brain–body interactions in recent years^17–24^, significant challenges remain in capturing the full complexity of interoceptive neural communication. Two-photon calcium imaging and fiber photometry have revealed the organization of sensory responses in subregions of the interoceptive system but are unable to monitor all interoception-relevant brain regions simultaneously. In contrast, whole-brain c-Fos mapping techniques provide a comprehensive overview of brain-wide activation patterns in response to interoceptive signals but do not report temporal dynamics. Thus, fully comprehending the brain-organ communication at the whole-brain scale – with single-cell resolution, fine temporal precision, during behavior – remains elusive. Compounding these limitations, the size and opacity of most animal bodies make it challenging to access ganglia of the peripheral nervous system, and it similarly limits access to the brain.

We set out to create a system that enables access to the dynamics of individual neurons across the entire brain and nodose ganglion – which relays signals from multiple organs to the brain – during delivery of nutrients to the gut in awake, behaving vertebrates. We aimed for a minimally invasive, all-optical system that delivers purely chemical stimulation to the gut without causing additional mechanical deformations or disruptions of normal gut physiology and behavior arising from invasive surgical methods or anaesthetics. We achieved this by delivering chemically caged compounds (such as glucose) by microgavage prior to experiments, using spatiotemporally precise photo-uncaging to release them within the gut during whole-brain imaging, and developing analysis algorithms for multimodal data. Since a major frontier in neuroscience is to unify systems neuroscience and the study of cognitive processes with the study of interoception and organ physiology, we incorporated visual stimulation and behavior readout into the system to enable controlled changes in interoceptive signals as animals behave and also receive exteroceptive input, i.e., input from the environment. Doing so allows for the simultaneous study of interoception, exteroception, and behavior at the whole-brain scale, at single-cell resolution, and without the need for surgery, all in a vertebrate animal. Since much of the central and especially the peripheral nervous systems are conserved across many vertebrate species, including mammals, this all-optical technique, combined with whole-brain analysis methods developed here, promises to reveal fundamental principles of nervous system function across vertebrate brains that have not previously been accessible.

## Results

We used a whole-brain light-sheet imaging system^25^ and extended it to have the ability to optically uncage chemicals in the body of larval zebrafish. This optical uncaging occurred through a targeted ultraviolet (UV) beam (30 μm-diameter, 355 nm-wavelength) that was routed through one of the horizontal illumination objectives and could be steered with a pair of galvo mirrors (**Extended Data Fig. 1**). This system can perform calcium imaging during nutrient release in the gut in transgenic larval zebrafish *Tg(elavl3:H2B-GCaMP7f)*^26^ that express GCaMP7f in almost all neurons, enabling single-cell-resolution imaging of the entire brain at multiple volumes per second while delivering physiologically-meaningful compounds to specific body regions, including different parts of the gut (**Fig. 1A,B**). Prior to experimentation, we introduced one of several caged compounds^27–29^ into the zebrafish gut using microgavage^30^ (**Extended Data Fig. 2A, Methods**). The transparency of larval zebrafish allowed us to then optically uncage the compound with the UV beam targeted to the gut, releasing the nutrient during whole-brain imaging while maintaining precise spatial and temporal control over gut stimulation in awake and behaving zebrafish, with no imaging or behavioral artifacts that might arise from mechanical methods of nutrient delivery (**Fig. 1B, Extended Data Fig. 2B,C**). To disambiguate neural encoding of gut-interoceptive signals from visual responses that might be elicited by scattered UV light, we also targeted the beam to a region outside the gut (either outside the fish or on the tail or swim bladder) on control trials. The setup also contained fictive-swim recording electrodes, a visual display under the fish, and closed-loop control over the visual stimulus by behavior to provide a virtual reality environment for the fish^25^. This setup allowed us to simultaneously interrogate interoceptive and exteroceptive processes and behavior.

**Figure 1.**
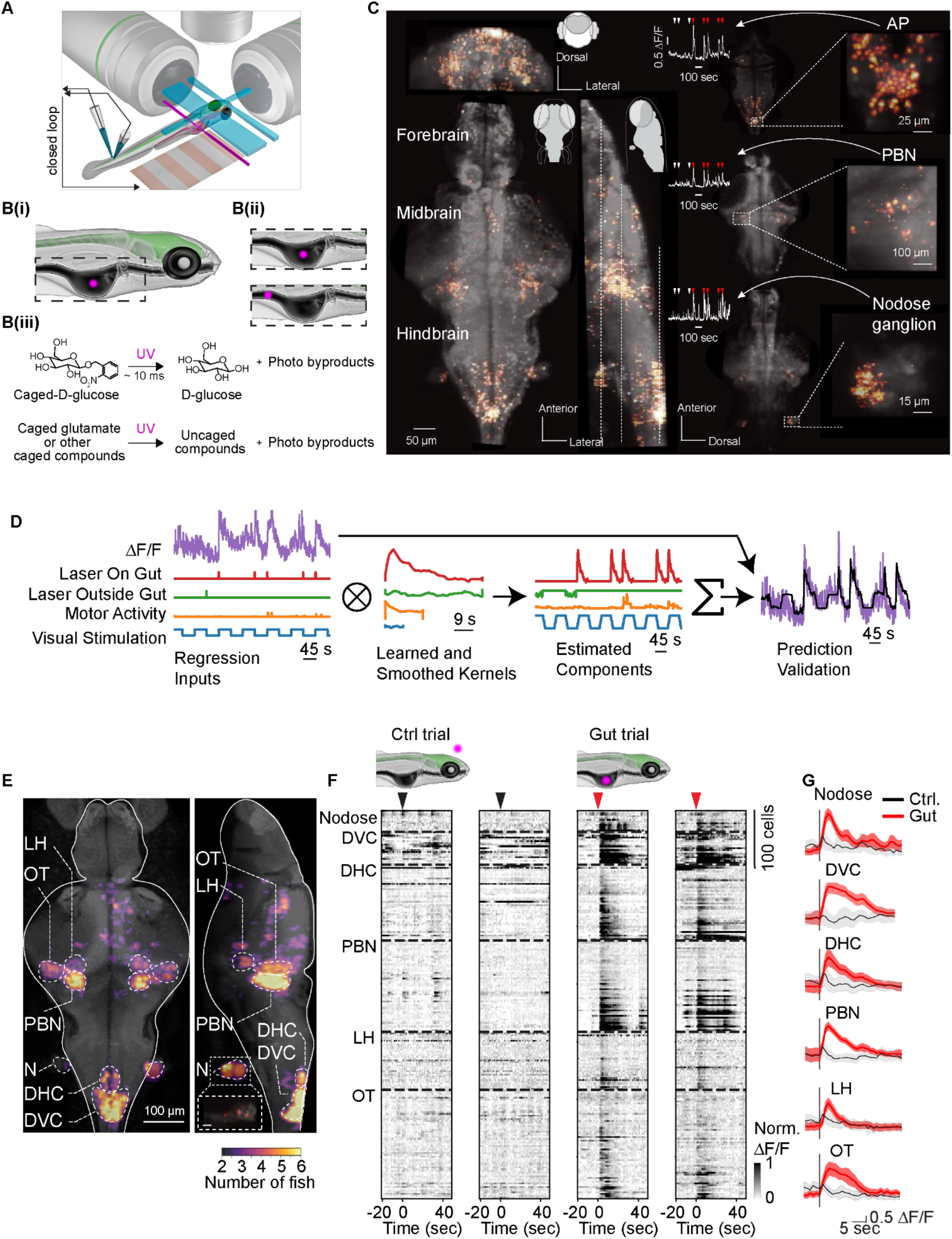
Whole-brain light-sheet imaging with integrated uncaging for all-optical nutrient delivery to the gut. **(A)** Diagram of the custom light-sheet microscope with integrated optical unaging. The blue planes denote the excitation light-sheets, and the purple beam represents UV illumination for uncaging. The central nervous system of a larval *Tg(elavl3:H2B-GCaMP7f)* zebrafish is depicted in green. This setup also enables real-time observation of larval behavior in a virtual reality environment, where fictive swimming signals (from motor neuron axon recording) drive instantaneous visual feedback throughout the experiment. **(B)** Noninvasive, rapid, and region-specific optical uncaging of caged chemicals in the larval zebrafish gut. B(i,ii): Representative cartoon showing optic uncaging and zoomed-in views to show localized nutrient release at fore- and midgut. Purple dot indicates UV beam. B(iii): The optical uncaging reaction yields free glucose within ∼10 ms. **(C)** Example of brain-wide average ΔF/F over 5 trials during glucose uncaging, shown in coronal, sagittal, and transverse views. Insets highlight example activity traces and images from the AP, PBN, and nodose regions. Activity maps were generated from light-sheet calcium imaging of neuronal responses, with fluorescence changes indicating activation across brain regions. Red arrowheads mark stimulus delivery times. See also **Suppl. Video 1.** **(D)** Lagged fused ridge regression for identifying stimulus sensitivities in neurons. A neuron’s activity (purple, left) is decomposed into components based upon regressors (left) by identifying component linear kernels (middle) that accurately predict a cell’s activity. Predictions are regularized (right) in both amplitude and smoothness. See Methods for further details. **(E)** Average brain-wide activity map in response to glucose uncaging in the gut (N = 6 fish). Color indicates the number of fish in which each region showed significant activation. Broad nodose activation in the combined map reflects inter-individual and registration variability due to increased error from reduced imaging clarity at greater depths. Inset shows an individual-animal example (see Methods). N: Nodose, DVC: dorsal vagal complex, PBN: parabrachial nucleus, LH: lateral hypothalamus, OT: optic tectum, DHC: dorsomedial hindbrain cluster. Insert is a zoomed-in view of nodose. Scale bar is 10 µm. **(F)** Example raster plots of neuronal activity during control and gut-targeted UV stimulation. Each column corresponds to a single trial. Cartoons depict the UV location, with dots for control (delivered outside the fish) and gut-targeted trials (dots are enlarged for illustration purpose). Each row corresponds to one neuron, grouped by brain region, with dashed horizontal lines separating neurons from distinct brain regions, black arrows indicating the delivery of control pulses, and red arrows indicating delivery of gut pulses. Time = 0 s marks the onset of UV stimulation. Cells were sorted within brain region by response duration. **(G)** Example average ΔF/F calcium traces (mean ± SEM) from control (gray) versus gut-stimulated (red) trials in representative brain areas.

Uncaging of caged D-glucose, a compound naturally produced by digestion of longer carbohydrates in the larval zebrafish gut and present in food sources^31^, elicited strong, time-locked responses in neurons in multiple regions throughout the peripheral nervous system and brain, including the nodose ganglia, parabrachial nucleus (PBN), and area postrema (AP) (**Fig. 1C; Suppl. Video 1**). To quantify neuronal responses to gut stimulation and assess their consistency across animals, we developed a regression model that predicts single-neuron activity based on time series gut stimuli, visual input, and behavior (**Fig. 1D, Methods**). Rather than *a priori* defining a single kernel shape for all cells globally, we utilize lagged regressors to simultaneously estimate linear response kernels for each regressor modality and each individual cell independently. This allows for differences in response kinetics both between cells as well as between regressors, improving model predictive accuracy by fine-tuning each individual cell’s response profiles. Importantly, the kernels are constructed in a fashion allowing for tractable close-form solutions (**Methods**, equation 2), enabling efficient computation even with hundreds of thousands of cell time series and hours of recording time. This method, then, can quickly estimate and decompose the contributions of gut stimuli, control UV light, visual stimuli, and motor swimming signals to the activity of every neuron.

Recording from a cohort of fish and applying our analysis methods, we created a whole-brain map (**Methods**) of gut interoception (also known as enteroception) showing the anatomical location of cells identified as gut-sensitive (**Fig. 1E**), which can be decomposed into individual activity traces (**Fig. 1F**) and the regional averages (**Fig. 1G**). This analysis revealed multiple regions with responses time locked to gut chemosensory events, including the optic tectum (OT), PBN, nodose ganglion (N), lateral hypothalamus (LH), a dorsomedial hindbrain cluster (which we term DHC), and dorsal vagal complex (DVC) consisting of the AP, the nucleus of the solitary tract (NTS), and the dorsal motor nucleus of the vagus (DMV). Thus, the methodology establishes a vertebrate whole-brain map of neuronal dynamics underlying gut interoception at single-cell resolution. Notably, our whole-brain imaging system allows for all these regions and their dynamics to be efficiently and reliably identified in a single experiment (**Fig. 1C**), rather than requiring multiple experiments focused on single brain areas.

The platform can be used to study whole-brain neuronal representations of a variety of nutrients in the digestive system, making use of advances in chemistry to photo-cage a wide variety of compounds^27–29,32–35^. Previously, we examined neural responses to caged glucose and created a whole-brain response map (**Figs. 1E, 2A**). Next, we used caged glutamate, an amino acid that is present in food^31^, and created a whole-brain response map (**Fig. 2B**). Although fine-scale differences appeared to be present, the brain areas responding to glutamate uncaging in the gut overlapped heavily with those responding to glucose, suggesting that population codes with these brain regions code for chemical identity but that there is no chemical-specific specialization of brain regions within the tested set of nutrients.

**Figure 2.**
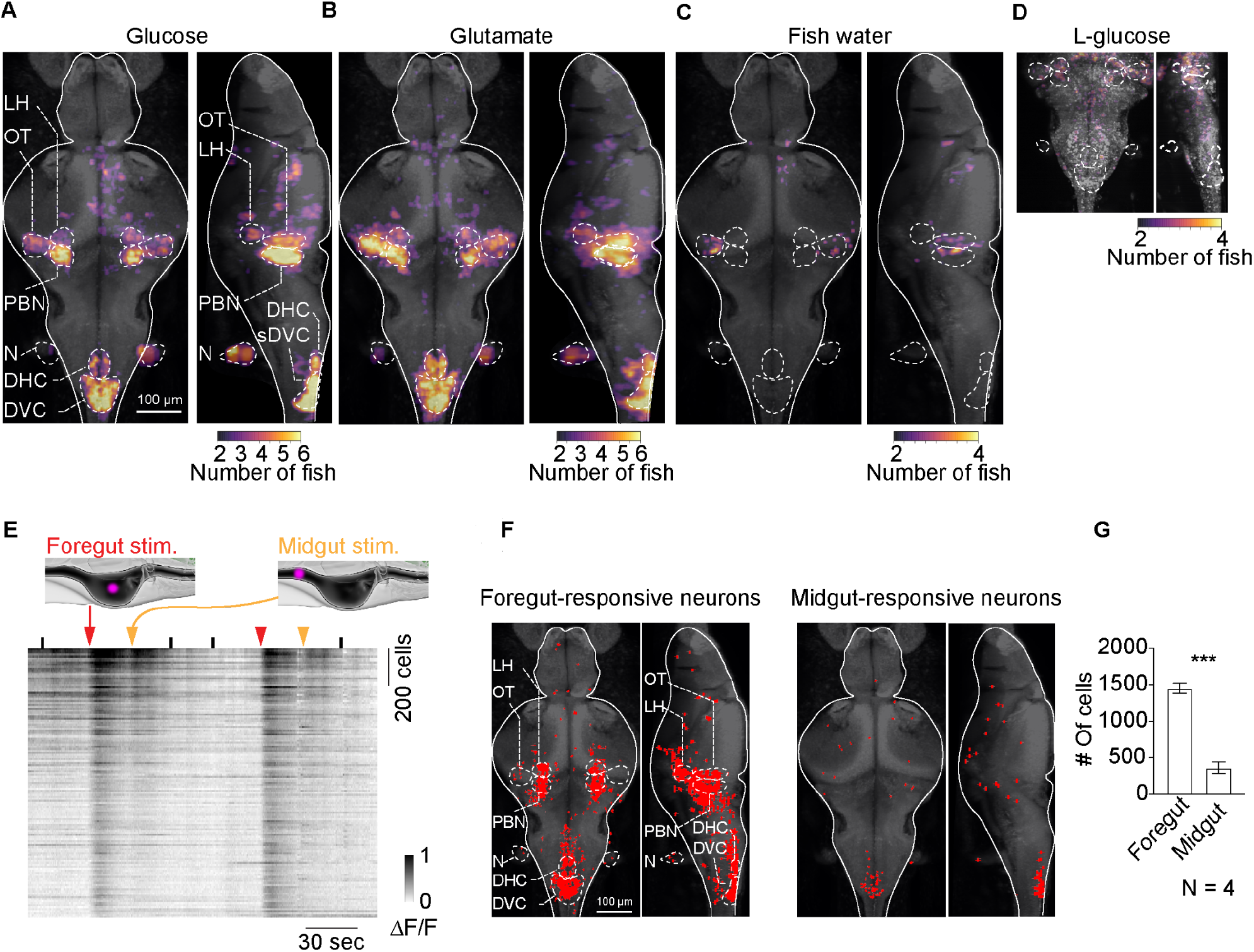
Chemical and spatial topography of neuronal responses to nutrients. **(A)** Average brain-wide activity map in response to glucose uncaging in the gut (N = 6 fish). Color indicates the number of fish in which each region showed significant activation. Same as Fig. 1E. Repeated here for easier comparison with (B) and (C). **(B)** Average brain-wide activity map in response to glutamate uncaging in the gut (N = 6 fish). **(C)** Control experiment showing the average brain-wide activity map in response to UV stimulation in the gut when only fish water is delivered, rather than a caged compound (N = 4 fish). This controls for potential artifacts such as tissue heating or other UV-evoked sensation. **(D)** Control experiment showing the average activity map in response to UV illumination in the gut with L-glucose (N = 4 fish). Only the section of the brain containing brain regions previously identified as consistently activated during gut stimulation was imaged. **(E)** Example raster plot of neuronal activity during foregut and midgut stimulation. Diagrams showing the UV location, with dots for UV stimulation on the foregut and the midgut. Each row represents a single neuron. Black tick marks the onset of control UV stimulation, where UV was targeted outside of the fish. Red arrowhead marks the onset of UV stimulation on the foregut. Yellow arrowhead marks the onset of UV stimulation of the midgut. Cells were sorted by max intensity. **(F)** Example brain maps of brain-wide foregut-responsive (left) versus midgut-responsive (right) neurons from one fish. Activated cells are coded in red. **(G)** Summary data showing the number of activated neurons for foregut versus midgut stimulation (N = 4 fish); bar graph represents mean ± SEM. ***p < 0.001.

Although we included control trials during which the UV localized outside the gut, we sought to apply a second set of control experiments to rule out artifacts of the UV light on brain responses. We replaced the caged compound solution with fish water as an additional control for UV artifacts, and found a negligible number of responding cells (**Fig. 2C**). We next verified that the chemical cages released as a byproduct of glutamate and glucose uncaging could likely not be detected by the gut, performing across-fish controls by replacing caged D-glucose with L-glucose (**Extended Data Fig. 3A-I)**, a metabolically inactive enantiomer of D-glucose that releases the same byproducts during uncaging. Uncaging L-glucose didn’t activate the brain regions activated during D-glucose or glutamate stimulation (**Fig. 2D**), showing both that the gut is specialized to sense D-glucose over L-glucose, and showing that the chemical cages elicit little or no responses and may be undetected by the gut epithelium. These additional control experiments provide validation for the accuracy and specificity of the paradigm. Notably, multiple compounds can be introduced to the same fish consecutively (**Extended Data Fig. 3J**), in principle enabling the creation of a cell-by-cell map across the entire brain to define the gut-chemosensory receptive field of individual neurons.

Due to optical access to the entire body, the optical system can be used to probe the brain’s representation of nutrient presence in specific locations of the digestive tract. We photo-uncaged glucose in the foregut and midgut in the same animal and observed that foregut signals led to strong brain-wide responses, while midgut stimulation led to substantially weaker responses (**Fig. 2E-G**). Thus, all-optical studies of interoception can be used to probe the topographic relationship between the location within organs and brain activity, showing, in this case, that the brain is able to sense the contents of the foregut more reliably than the contents of the midgut.

In natural settings, rather than processing internal signals independently, the brain combines information about the body’s nutrient state with external sensory cues to dynamically shape behavior^36–42^. To understand how the brain integrates interoceptive information with exteroceptive and motor signals, we next examined the interaction between gut stimulation, visual processing, and behavior. Here, we presented drifting grating stimuli to the zebrafish to elicit optomotor responses while recording both motor activity and brain responses. During gut stimulation, zebrafish continued to display standard optomotor responses, and visual and motor maps created during gut stimulation experiments resembled previous findings^43,44^ (**Extended Data Fig. 4**), indicating that our paradigm preserves visuo-motor processing and behavior.

We sought to assess whether or not – and to what degree – a gut sensitive neuron’s activity could be better explained when incorporating visual stimulation and motor activity. To do so, we assessed the predictive benefits of linear models (based on those of **Fig. 1D**) incorporating all regressor modalities (gut and control UV stimulation, visual stimulation, and motor activity) over those containing only UV gut and control regressors. A subset of gut-sensitive neurons responded to both gut stimulation and motor activity, and another responded to a combination of gut stimulation, motor activity, and visual stimulation (**Fig. 3A,B**). In total, 26% of cells across six fish showed improvements in the correlation between real and predicted responses of 0.1 or greater when visual and motor information was included in models of neural activity compared to models that were only based on gut information (**Fig. 3C**). We attempted to further identify which modalities most strongly affect responses in gut-sensitive neurons – and the spatial patterns of multimodal neurons – by computing the average increase in performance when incorporating either motor activity (GM, incorporating gut and motor signals), visual stimulation (GV, incorporating gut and visual signals), or both (GMV, incorporating all three signals) as a function of brain area (**Fig. 3D,E**), as well as raw correlations between neural activity and visual stimulation (**Fig. 3F**) and motor activity (**Fig. 3G**). Throughout the brain, many gut-sensitive neurons also exhibited motor and visual responses, with gut-sensitive areas within the posterior brainstem also strongly motor sensitive and gut-sensitive areas within the midbrain responsive to both motor activity and visual stimulation.

**Figure 3.**
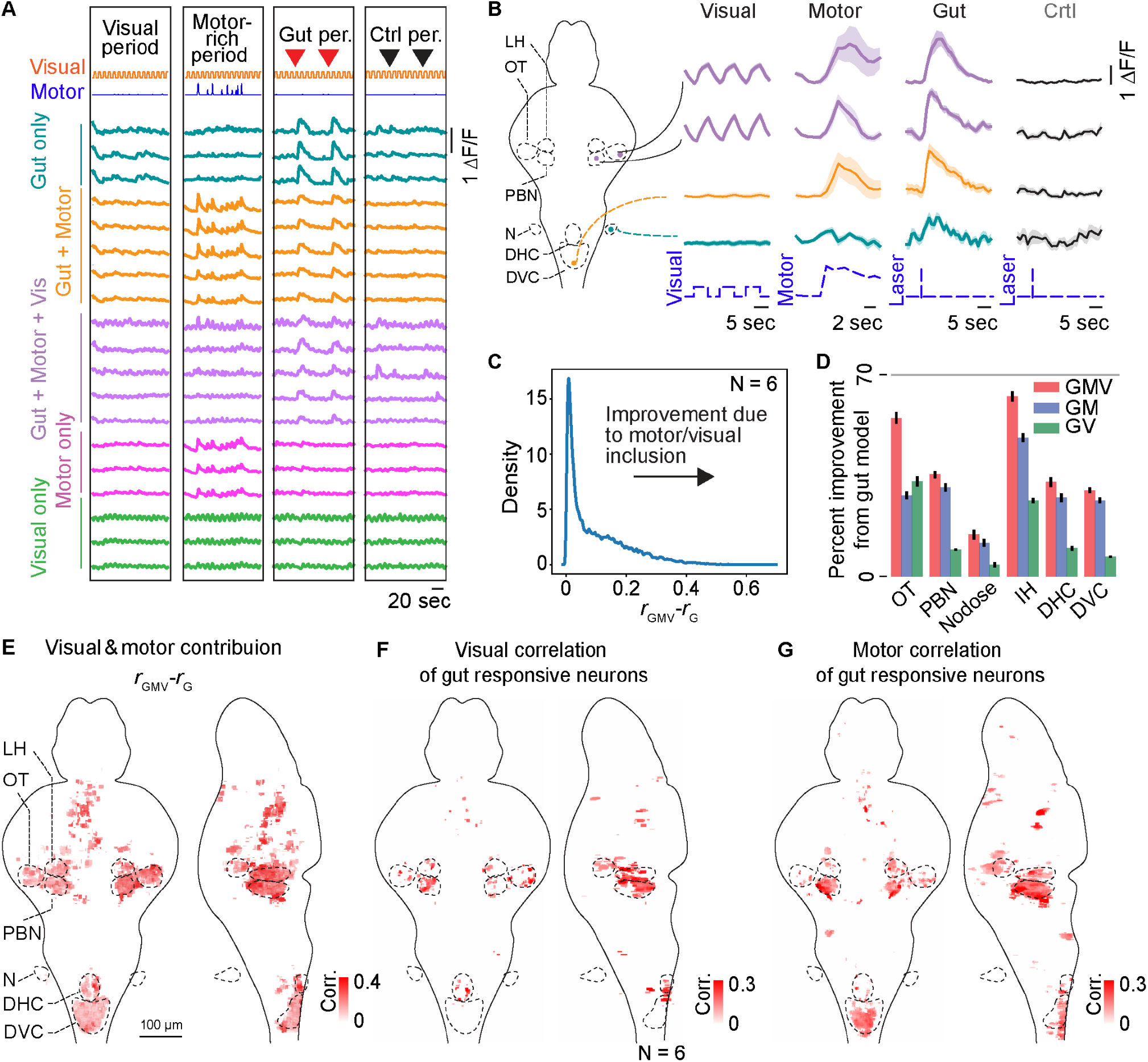
Integration of sensory, motor, and gut signals across the brain. **(A)** Example calcium traces from individual gut-responsive neurons during four distinct experimental periods (visual, motor, control, and gut). Each row represents a single neuron. The top orange trace indicates visual motion, with upward segments representing forward movement of the visual gratings and downward segments indicating no motion. The blue trace reflects motor power derived from a sliding window of the tail voltage signal. The red arrowheads mark the onset of each UV stimulation on the foregut. The black arrowheads mark the onset of control UV stimulation just outside the fish. **(B)** Example average ΔF/F responses of representative single neurons from major gut-responsive brain regions. Each dot on the schematic (left) marks the location of a sampled neuron. Plotted are the median and SEM. The dashed lines at the bottom mark the timing of visual, motor, gut, and control stimulation. **(C)** Distribution of *r*_*GMV*_ − *r*_*G*_, the difference in correlation between a neuron’s activity and a linear model incorporating gut, motor, and visual information and the correlation between the neuron and a linear model only incorporating gut information. Note the strong right tail, indicating the presence of multimodal neurons. **(D)** Bar plots of percent change in correlation from the gut model and a model including gut, motor, and visual information (red), a model including gut and motor information (blue), and a model including gut and visual information (green), separated by brain region. Note that regions found in the hindbrain have minimal improvement when including visual information. G,M,V stands for gut, motor, visual, so that, for instance, GMV represents a model based on gut, motor, and visual signals, and GM represents a model based on gut and motor signals. **(E)** Spatial map of *r*_*GMV*_ − *r*_*G*_ across the brain (N=6 fish). Increased color saturation indicates additional predictability gained when incorporating motor and visual information in models of neural activity. **(F)** Spatial map indicating the correlation of gut-responsive neurons with the visual signal across the brain (N = 6 fish). The color scale indicates the correlation coefficient between neuronal activity and visual motion. (See Methods for analysis details.) **(G)** Spatial map of the same gut-responsive neurons showing correlations with the motor signal (N = 6 fish). Color indicates the correlation coefficient between neuronal activity and motor power. (See Methods for analysis details.)

As our system images the whole brain nearly simultaneously, we have the additional benefit of being able to observe trial-to-trial correlations between gut-responsive neurons in disparate regions (**Fig. 4A**, see **Extended Data Fig. 5A** for additional traces). We found that the signal correlation^45^ (the correlation between trial-average gut responses across identified cells, **Methods**) between gut-responsive neurons was high across gut-responsive brain areas, even across distant regions (**Fig. 4B**). Noise correlations (correlations in trial-to-trial variations between cells, Methods), are commonly used in neuroscience to quantify the degree of functional connectivity between pairs of neurons; correlated fluctuations present in two neurons could arise from either spontaneous events occurring in common inputs to both neurons or spontaneous events in one neuron impacting the other. We quantified average noise correlations within pairs of neurons across brain regions, observing heightened values between cells within DVC, parabrachial nucleus, and the DHC compared to control populations (non-gut coding cells in the nodose ganglia) (**Fig. 4C**). These quantifications suggest the existence of a tightly coupled functional network for interoceptive signaling across the brain.

**Figure 4.**
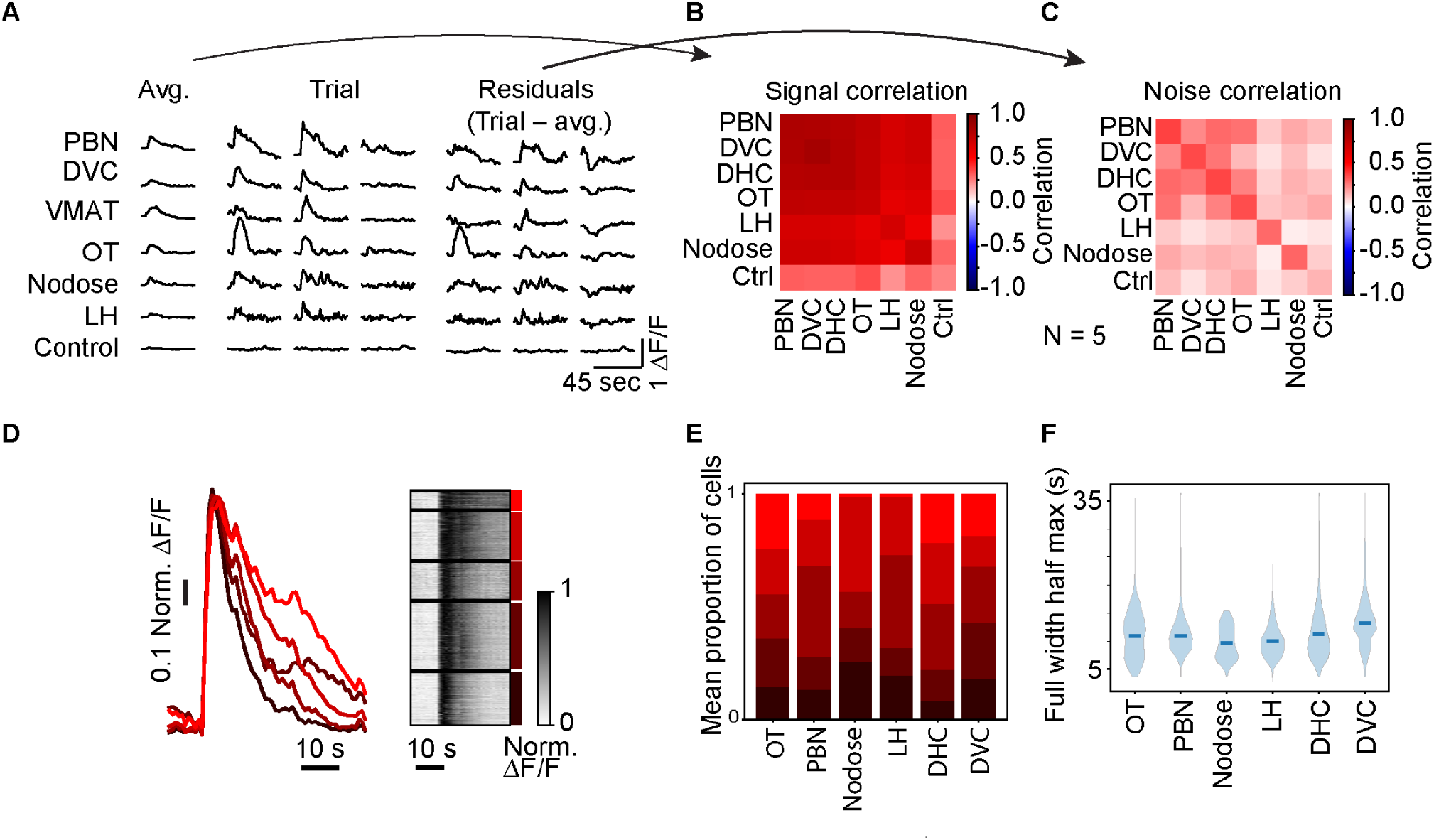
Correlations and brain-wide dynamics of gut-responsive neurons. **(A)** Example average (left), trial responses (middle), and residuals (right) for example gut-sensitive neurons in identified areas as well as control. Each trace is approximately 55 s long. **(B)** Mean signal correlation between gut-sensitive neurons in identified regions. Note the strong signal correlations present between brain regions, indicating similar representations of gut stimulation across the brain. **(C)** Mean noise correlation. Note that areas within the brain show high noise correlations between one another, as opposed to nodose or control. **(D)** Results of clustering average responses to gut stimulation for all gut-sensitive neurons identified in a single fish. *Left:* Mean responses within each cluster demonstrate the variability in decay time found across the brain. *Right:* Raster plot of all responses grouped by cluster (as in left). **(E)** Mean proportion of cells found in each cluster for identified gut-sensitive areas. Clusters are colored by ordering decay timescales, as in (C). Note that all areas contain all different response types with varying frequency. **(F)** Distribution of full-width half-maxes in each region.

We next inspected the response time courses across the brain. While the responses of most neurons showed an onset time of five seconds or less, their decay time varied from ∼10 seconds to over a minute. To illustrate this diversity in decay times, we used k-means clustering to divide the average responses for each fish into five clusters, resulting in clusters segregated largely on the basis of decay time. By performing this analysis on each fish individually and then sorting the clusters by decay time, we could compare how decay times vary across the fish brain (**Fig. 4D**). We initially hypothesized that these varying time courses would be due to region-specific properties, with peripheral regions like the nodose exhibiting faster responses and downstream regions like the PBN exhibiting more integrative and sustained responses. We instead found that all regions contained neurons tiling most of the temporal spectrum, with each brain region containing multiple different response types (**Fig. 4E**). Despite the broad distributions of timescales contained within each group (**Fig. 4F**), some brain regions nonetheless showed significant differences (**Extended Data Fig. 5B**). Notably, the nodose ganglia had a significantly shorter response than most other detected brain regions, consistent with this being a peripheral sensory locus of the vagal interoceptive nervous system between the gut and the brain. While it is possible that this is a readout of the lingering uncaged chemical in the gut, it may indicate, consistent with the above analysis of inter-region correlation, that these intermingled timescales are due to across-area communication.

In sum, we developed a system for whole-brain interrogation of gut sensation in awake and behaving larval zebrafish. This presents a new paradigm for merging analyses of interoceptive and exteroceptive processing, as the system allows for holistically combining both stimulation modalities – as well as behavior – into a single framework. This work enables the investigation of the neural basis of complex behaviors that rely on interoceptive and exteroceptive signals, unifying systems neuroscience with the study of brain-body communication.

## Discussion

The system developed in this work was applied to the study of gut-brain communication but, as a general paradigm for whole-brain imaging during chemical manipulation of bodily tissue, is widely applicable to the study of brain-body interactions. The vast array of existing caged chemicals^27–29,32–35^ implies that the same paradigm can be used for studies of the brain and behavioral response to many other important perturbations of body physiology. Within the gut, by systematically varying the uncaged compounds, future studies will dissect how the brain encodes specific nutrient classes, detects harmful substances, and integrates metabolic signals into feeding and homeostatic regulation. By making use of the zebrafish’s ability to absorb many small molecules through the bath or injecting caged compounds directly into the bloodstream or specific tissues, applications beyond the gut include the study of neural encoding of light-induced liver acidification^46,47^, blood glucose level changes^48,49^, hormones in the bloodstream^50,51^, optogenetically-induced apoptosis in various body organs^52^, and more. These future applications promise new insights into the brain’s encoding of many body-wide physiological signals, as well as their influence on perception, memory, and behavior, that were hitherto inaccessible.

The ability to monitor such body-brain interactions at the whole-brain level at single-cell resolution enables precise dissection of the mechanisms by which internal physiological states modulate perception and action. In this work, by embedding gut stimulation within a virtual reality and whole-brain imaging setup, we provided a noninvasive way to modulate body physiology while simultaneously tracking behavior and imaging neural responses across the entire brain, allowing us to examine how gut-derived signals influence ongoing motor and sensory processing. The influence of bodily state on brain-wide processing is profound, as exemplified by the impact of sickness or hunger on behavior, emotional state, and as reinforcement signals in learning^13–15^. These effects are only rarely incorporated into systems-neuroscience studies, in part because accessing the brain and body simultaneously has been challenging, and in part because the sites of interaction are often unknown. Whole-brain imaging during body manipulation overcomes these challenges, allowing for the monitoring of interactions across brain areas implementing intero- and exteroceptive computations – and beyond the virtual reality environments used here, other scenarios, including the influences of body physiology on perception and action in prey capture and learning. Eventually, through the lens of control theory, the merging of systems neuroscience with studies of interoception promise to lead to an understanding of the body-wide implementation of control algorithms jointly governing motor actions and physiological control.

The observation that many gut-responsive neurons also react to visual or motor signals suggests a more integrative role for these neurons in coordinating behavior than simply transmitting interoceptive cues. This multimodal responsiveness implies that the brain does not process internal signals in isolation; instead, it integrates information about the body’s nutrient state with external sensory inputs to guide behavior adaptively. This finding aligns with the broader framework in behavioral neuroscience, where internal states influence not only neuronal activity but also interact with external signals to modify complex behaviors. By bridging the gap between interoception and exteroception, these multi-sensory gut-responsive neurons could serve as critical nodes in the circuitry ensuring that the animal’s behavior is finely tuned to both its internal needs and its external environment.

Our paradigm also provides opportunities for studying the developmental time course of interoception. The fish’s small size and transparency permit whole-brain imaging in larval and juvenile zebrafish for the first several weeks of life. This presents the opportunity to investigate how gut innervation, gut-brain functional interactions, and the full repertoire of gut-related behaviors co-develop from hatching to maturity^53^. Applying our general paradigm to *Danionella cerebrum* would allow for the study of brain mechanisms of interoception in adults, as its small size and transparency persist into adulthood^54^. This allows for investigations of how interoceptive signals affect more complex cognitive behaviors that first appear in adult fish, including a variety of social behaviors^55^ and sophisticated learning and memory tasks^56^.

Overall, this platform enables investigations of interoceptive processing at an unprecedented level of detail. It opens new opportunities for studying how internal bodily signals interact with sensory and motor circuits, how the brain maintains homeostasis, and how physiological states shape behavior. By offering a noninvasive, high-resolution, and versatile approach to manipulating and monitoring interoceptive signals, our system sets the stage for future research into the neural basis of brain-body communication.

## Acknowledgements

We thank Xuesong Li, and Kaspar Podgorski for their advice on the UV lightpath, Vasily Goncharov for building the customized attenuator for the UV laser, Alex Sohn for customizing and building imaging chambers and chamber holders, Igor Siwanowicz for making the 3D anatomy schematics, Jeff Talbot and Jon Arnold for making the 3D illustration of our light-sheet and UV lightpath, Janelia Vivarium for the support, Hari Shroff, Eric Schreiter, Carsen Stringer, Yin Liu, Anoj Ilanges, Isabel Espinosa Medina, Dhruv Zocchi, and Emmanuel Marquez Legorreta for feedback on the manuscript, and all Ahrens and Fitzgerald lab members for feedback throughout the project. This work was supported by the Howard Hughes Medical Institute. JEF acknowledges support from the National Institute for Theory and Mathematics in Biology through the National Science Foundation (grant number DMS-2235451) and the Simons Foundation (grant number MPTMPS-00005320).

## Methods

### Key resources table

| REAGENT or RESOURCE | SOURCE | IDENTIFIER |
| --- | --- | --- |
| **Chemicals, peptides, and recombinant proteins** |  |  |
| Caged D-glucose | This paper |  |
| Caged L-glucose | This paper |  |
| MNI-caged L-glutamate | Tocris | Cat# 1490 |
| Tricaine | Sigma-Aldrich | Cat# A5040 |
| methylcellulose | Sigma-Aldrich | Cat# M 0387 |
| **Experimental models: Organisms/Strains** |  |  |
| Tg(elavl3:H2B-GCaMP7f) | Yang et al.<sup>26</sup> | jf90 |
| **Software and algorithms** |  |  |
| FIJI | NIH Image | https://imagej.nih.gov/ij/ |
| Adobe Illustrator | Adobe, San Jose, CA | https://www.adobe.com/products/illustrator.html |
| Custom code | This paper | https://github.com/benJames1212/gutBrainPipeline |
| Segmentation software | Mu et al.<sup>57</sup> | https://github.com/mikarubi/voluseg |

### Zebrafish

Larvae were reared at 28.5°C under a 14:10-hour light-dark cycle. Rotifers were provided as food during the rearing period. Zebrafish larvae aged 7–8 days post-fertilization (dpf) were selected for experiments. The transgenic line *Tg(elavl3:H2B-jGCaMP7f)*^jf90^ was used for all experiments. All experimental procedures were conducted in compliance with institutional animal care guidelines and were approved by the Institutional Care and Use Committee of Janelia Research Campus (JRC).

### UV–Vis Spectroscopy

Spectroscopy was performed using 1-cm path length, 3.5-mL quartz cuvettes from Starna Cells or 1-cm path length, 1.0-mL quartz microcuvettes from Hellma. All measurements were taken at ambient temperature (22 ± 2 °C). Absorption spectra were recorded on a Cary Model 100 spectrometer (Agilent). Samples (50 μM) were prepared in phosphate-buffered saline (PBS), pH 7.4 for measurements.

### Analysis of photolysis products by tandem high-pressure liquid chromatography–mass spectrometry (LC–MS)

To analyze photolysis products, solutions of caged-D-glucose or caged-L-glucose (200 μM) were prepared in PBS and placed in a glass vial. An aliquot of this freshly prepared solution was immediately analyzed by tandem high-pressure liquid chromatography–mass spectrometry (LC– MS) using an Agilent 1200 LC–MS system equipped with an autosampler, diode array detector, and mass spectrometry detector (ESI; positive ion mode) using a 4.6 × 150 mm Gemini NX-C18 column with a 5–95% or 5–50% gradient of CH3CN in H2O containing constant 0.1% (v/v) TFA. Chromatograms were monitored using absorbance at 254 nm. The solution was then irradiated with 405 nm light from an LED array (LOCTITE CL20 flood array) followed by analysis using LC– MS.

### Synthesis and characterization of Caged-D-glucose and Caged-L-glucose

Caged-D-glucose was synthesized as previously described^28^. The enantiomer, caged-L-glucose, was synthesized using the same protocol but using L-glucose as the starting material. As expected, the enantiomeric pair of caged-D-glucose and caged-L-glucose show effectively identical 1H NMR spectra, 13C NMR spectra, absorption spectra, and photolysis traces (**Extended Data Fig. 3**). Caged-L-glucose: 1H NMR (CD3OD, 400 MHz) δ 8.07 (dd, J = 8.1, 1.3 Hz, 1H), 8.02 (dd, J = 7.9, 1.3 Hz, 1H), 7.71 (td, J = 7.6, 1.3 Hz, 1H), 7.56 – 7.46 (m, 1H), 5.28 and 5.08 (AB, JAB = 15.2 Hz, 1H), 4.43 (d, J = 7.6 Hz, 1H), 3.88 (dd, J = 11.9, 1.9 Hz, 1H), 3.68 (dd, J = 11.9, 5.2 Hz, 1H), 3.41 – 3.31 (m, 4H). 13C NMR (CD3OD, 101 MHz) δ 148.8 (C), 135.7 (C), 134.7 (CH), 130.3 (CH), 129.2 (CH), 125.5 (CH), 104.0 (CH), 78.1 (CH), 78.1 (CH), 75.2 (CH), 71.6 (CH), 68.4 (CH2), 62.7 (CH2). MS (ESI) calcd for C13H19NO9[M+H2O]+ 333.1, found 333.0. To confirm these two compounds are enantiomers the specific optical rotation of caged-L-glucose was measured to be [α]21D = 35±1° (c =0.1, pyridine). This is opposite of the literature value of caged-D-glucose, which was measured to be [α]21D= −36.4° (c =0.55, pyridine).

### Uncaging light path

We customized the UV light path on our light-sheet microscope to enable precise uncaging of compounds. The UV light source was a 7W max, 355 nm pulsed laser with repetition rate of 2 kHz (3502-100, DPSS Laser Inc.). To control the UV intensity, a custom beam attenuator, consisting of half-wave plate (WPHSM05-355, Thorlabs) mounted on a rotation mount (CRM05, Thorlabs), High Power Laser-Line polarized beam splitter (CCM1-PBS25-355-HP, Thorlabs), Beam Trap (BT600, Thorlabs), was attached directly to the laser head. The attenuated beam was directed onto a pair of galvo mirrors (1 & 2 in Extended Data Fig. 1, 6215K, Cambridge Technology), enabling independent horizontal and vertical positioning. The reflected beam was then directed through a scan-relay system (L1 in Extended Data Fig. 1, LA1509-ML, f = 100 mm & L2 in Extended Data Fig. 1, CA254-200-A-ML, f = 200 mm, Thorlabs) and focused onto the sample using a light-sheet objective (Obj 1 in Extended Data Fig. 1, 4×/0.28 NA, Olympus). To ensure precise UV beam positioning on the sample, a calibration pathway was implemented on the opposite side. This pathway included an opposite objective (Obj 4 in Extended Data Fig. 1, 4×/0.28 NA, Olympus), a lens (L4 in Extended Data Fig. 1, LA1509, Thorlabs), a shortpass filter (FESH0450, Thorlabs), and a camera (BFS-U3-32S4C-C, FLIR Integrated Imaging Solutions Inc.), enabling real-time visualization for UV beam alignment. This setup ensured accurate and reproducible UV beam delivery to the sample.

### Light-sheet microscopy

The light-sheet microscope and behavioral setup used in this study were designed as described previously^25^. Briefly, the detection pathway included a vertically mounted water-dipping objective (Obj3 in Extended Data Fig. 1, 16x/0.8 NA, Nikon) attached to a piezo stage (P-725.4CD PIFOC, Physik Instrumente), along with a tube lens (TL3 in Extended Data Fig. 1, f = 200 mm) and an sCMOS camera (Orca Flash 4.0, Hamamatsu). Fluorescence separation was achieved using a band-pass filter (525/50 nm, Semrock) for GCaMP signals, effectively isolating them from scattered laser light (488 nm). Both of the two illumination arms were equipped with galvanometer scanners (3-6 in Extended Data Fig. 1, 6210, Cambridge Technology), an f-theta lens (SL 1 & 2 in Extended Data Fig. 1, SL50-CLS2, f = 50 mm, Thorlabs), a tube lens (TL 1 & 2 in Extended Data Fig. 1, TTL200MP, f = 200 mm, Thorlabs), and an air illumination objective (Obj 1 & 2 in Extended Data Fig. 1, 4x/0.28 NA, Olympus). The galvanometer scanners and f-theta lens enabled lateral and axial scanning of the collimated laser beams (488 nm). To prevent direct retinal stimulation, the excitation laser was electronically switched off when it passed over the zebrafish eye.

### Microgavage

The microgavage method used in this study was adapted from previous work by Cocchiaro and Rawls^30^. In brief, zebrafish larvae were anesthetized using tricaine, embedded in 3% methylcellulose (Sigma-Aldrich), and a fine borosilicate glass capillaries (∼27–30 μm in diameter and blunt; BF100-58-10, Sutter Instrument) was used to deliver approximately 4.6 nL of the desired substance into the gut lumen under a dissecting microscope. Following the procedure, larvae were allowed to recover for 5–10 minutes until they resumed normal swimming behavior before the start of imaging experiments.

### Simultaneous optical uncaging in the gut and whole-brain light-sheet imaging

The preparation procedures followed previously established protocols^57^. Briefly, after recovering from microgavaging, non-paralyzed fish were embedded in 2% low-melting point agarose (Sigma-Aldrich) within a custom-made imaging chamber (CAD designs available upon request), agarose were carefully removed around the head to allow for optical access during imaging and from the tail for electrophysiological recordings.

The position, onset and duration of UV were controlled by custom software developed in C# (Microsoft) during simultaneous whole-brain light-sheet imaging. Each UV pulse consisted of smaller, 1 ms long pulses separated by 5 ms of off time for a total of 200 ms. The intensity of UV delivered to the gut is 0.7 mW.

For visuomotor transformation experiments, motor nerve signals were simultaneously recorded. Extracellular recordings of the motor nerve signal were obtained using borosilicate glass pipettes (TW150-3, World Precision Instruments) fabricated with a vertical puller (PC-10, Narishige) and polished using a microforge (MF-900, Narishige) to achieve a tip diameter of approximately 40 µm. Recordings were conducted in current-clamp mode using an Axon Multiclamp 700B amplifier, with signals acquired via National Instruments data acquisition boards. Data were sampled at 6 kHz and band-pass filtered with a low-pass cutoff at 3 kHz and a high-pass cutoff at 100 Hz.

Visual stimuli were also simultaneously delivered through custom software developed in C# (Microsoft). Visual projections were made from below the fish using a miniature projector (Sony Pico Mobile Projector, MP-CL1) connected to the data acquisition computer, with the image projected onto a diffusive plastic screen affixed to the outside of the chamber, approximately 1 cm beneath the fish. The experiments were conducted in one-dimensional virtual environments comprising regularly spaced red and black bars (∼2 mm thick) to enable imaging of GCaMP-expressing fish. The stripes moved forward at a constant velocity, regardless of the fish’s swimming behavior, to ensure consistent visual stimulation throughout the experiment.

### Segmentation and time series extraction

Cells were segmented and time series extracted using a previously described volumetric segmentation software^57^. In brief, brain volumes were first registered to an average volume, and spatial footprints for cell segments were initialized by identifying local fluorescence maxima within temporally correlated and spatially continuous voxels. We then used a constrained non-negative matrix factorization to approximately solve the following equation using alternating least squares: *V* ≈ *WH* + *XI*, where *V* is the (*n* × *t*) matrix of spatiotemporal fluorescence, *W* and *H* are the (*n* × *c*) and (*c* × *t*) matrices of spatial and temporal footprints of cell segments, and *X* and *I* are the (*n* x 1) and (1 × *t*) matrices of spatial and temporal traces of the background signal, and *n, c*, and *t* indicate voxels, cell segments, and time, respectively. Factorization of *W* and *W* was regularized to ensure spatial contiguity spatial and sparsity, as well as mean fluorescence amplitude. *Δ F*/*F* was then computed as: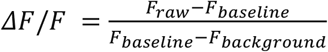, with *F*_*baseline*._ estimated from 10th lowest percentile of the raw fluorescence *F*_*raw*_ over a rolling 5-minute period, and *F*_*background*_ computed from voxels located within the imaging volume but outside the brain. Note that as the UV laser induces a large artifact in frames recorded during stimulation, any frames recorded during UV stimulation are removed and replaced with a copy of the previous frame.

### Construction of Brain Maps

Functional volumes were registered to the Z-Brain atlas^58^ using the BigStream software packages (https://github.com/JaneliaSciComp/bigstream). Following the registration of each fish onto a common brain, we first transformed the centroids of each identified gut-sensitive cell into a common anatomical space. We then expanded these centroids into cubes with side lengths of 5.5 µm in the xy axes and 6 µm in the z axis to minimize the effects of biological variability in brain anatomy and small errors in registration. In the nodose region, cubes measured 15 µm in the xy axes and 12 µm in the z axis to accommodate increased registration error resulting from reduced imaging clarity at greater depths. The resulting volume was then smoothed with a Guassian filter of standard deviation 1.624 µm.

### Regression analysis

In this work, regression fits were computed using fused distributed lag ridge regression, where ‘fused’ indicates a penalty on the magnitude of differences in temporally successive parameters^59^. As regressors, we used stimuli (visual drifting gratings and UV uncaging) and recorded motor activity. For a model taking a single time series, *x*, as input, a neuron’s response is modeled as 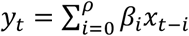, where *y* is a vector of neural activity of length *T* corresponding to the neuron’s activity across all *T* time points, *ρ* is the lag order that sets the length of time previous inputs are able to affect the current responses, and *s* is a vector of coefficients of length *ρ*. To extend this regression to allow for the analysis of multiple cells simultaneously, vectors are converted to matrices. In matrix notation, *Y* = *Xβ*, with (time × cells) matrix *Y*, (time × (*ρ*+1)) matrix *X*, and coefficient matrix *β* of size ((*ρ*+1) × cells). Note that the matrix *X* is constructed from a single time series, with each row shifted by one time point relative to the instantaneous time point.

We define a loss function:

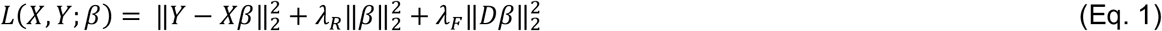

incorporating an L2 penalty on both coefficient magnitudes (second term on the RHS in Eq 1) and differences in temporally subsequent coefficient values to promote smoothness (third term on the RHS in Eq. 1, indexed by t). Here, *D* is a finite difference matrix (a matrix of size ((*ρ*+1) x (*ρ*+1)) with all diagonal entries equal to 2 and all one-off diagonal entries equal to negative one, see **Extended Data Fig. 6A**). Model parameters were selected to analytically minimize the loss function given in Eq. 1;

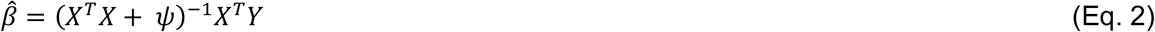

where *ψ* = *λ* _*R*_ *I* + *λ*_*F*_*D*^*T*^*D, I* is the identity matrix of size ((*ρ*+1) x (*ρ*+1)), *λ*_*R*_ is the usual ridge regularization strength, and *ρ*_*F*_ is the regularization strength promoting temporal smoothness in kernels. Values for both regularization parameters were set manually (*λ*_*F*_ = 2, *λ*_*F*_ = 20). For *ρ* > 0, the resulting vector β can be viewed as a response kernel, while for *ρ* = 0 the value of β is simply a coefficient.

Regressions taking multiple time series as input are performed analogously, with *X* = [*X*_1_, …, *X*_*N*_] now being of size (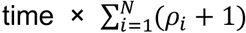) constructed by concatenating each regressor *X*_*i*_ with corresponding lag order *ρ*_*i*_, and β now being a matrix of size (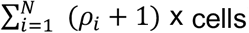). For this work, lag orders were set to generate kernels approximately 55 s for control and gut UV pulse responses, 10 s for motor responses, and 5 s for visual responses, chosen by identifying the point at which each corresponding response decayed back to baseline. As not all imaging frequencies are a multiple of 60 s, the exact number of frames used in the lag response varies from fish to fish. Note that when multiple regressors are used in a single analysis, the finite difference matrix *D* is modified to prevent penalties occurring between the boundaries of distinct regressors (see **Extended Data Fig. 6A**), with each element 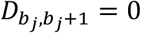 and 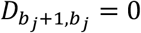 for each boundary index 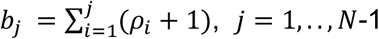. In other words, values of one-off diagonal elements of *D* occurring at the boundaries between regressors are set to zero, thereby ‘breaking’ the temporal fusion of regressors and preventing a penalty between the last time point of one regressor and the first of another regressor. All regressions other than those used in **Fig. 3** for estimating multi-modal activity were performed using leave-one-out cross-validation by leaving out periods of length 55 s beginning with each gut-targeted UV pulse. Regressions for estimating multi-modal activity were performed without cross-validation in order to utilize all data available. Regressors were zero-padded for *ρ* timesteps preceding neural activity in order to utilize all recorded experimental data.

### Gut-sensitive cell selection

Gut sensitive cells were identified using the fused lag ridge regression described above by creating a parts model. In this general paradigm, a ‘full’ model is first constructed utilizing all available regressor data and then compared to a ‘part’ model, which utilizes only a subset of the data incorporated in the ‘full’ model. For cell selection, the ‘full’ model consists of two regressors: the first corresponding to the time of gut-focused UV pulses, and the second corresponding to the time of *all* UV pulses (thus incorporating off-gut UV pulses as well as gut-focused UV pulses). The ‘part’ model incorporates only a single regressor corresponding to the timing of all UV pulses, identical to the second regressor in the ‘full’ model. Based on the results of the two models, cells are identified as responding to gut stimulation if they meet the following criteria: the full model yields significant correlations (see below), has a statistically better fit than the ‘parts’ model as assessed via a paired Wilcoxon signed-rank test between the difference in correlation between the ‘part’ and ‘full’ model for each gut-centered UV pulse, and the distribution of peak responses to gut pulses are statistically greater than the peak responses to control pulses, assessed via an unpaired Mann-Whitney test, with the last condition preventing false-positives in the form of miniature UV evoked responses.

### Assigning threshold for significant correlations

We assume we can divide the set of recorded neurons into two mutually exclusive groups: those responsive to gut stimulation, and those unresponsive to gut stimulation. As non-gut-responsive neurons by definition should not be predictable using gut stimulation regressors, the distribution of correlations between these neurons and predictive models utilizing only gut stimulation should be approximately normal with mean zero and variance set by noise. Gut-responsive neurons, on the other hand, should have a distribution of correlations centered at some value above zero, dependent upon the strength of a cell’s response to gut stimulation. Consequently, analysis of the distribution of correlations across all cells in the brain yields a distribution heavy-tailed on the positive side corresponding to gut-sensitive responses, while the negative side should be reflective of noise (**Extended Data Fig. 6B**). We thus constructed a null model by estimating the standard deviation, *σ*_*G*_, of the left hand side of a Gaussian distribution centered on the mode, *m*, of the data. We then utilized the parameters of this null model to set the threshold for significant correlations as 3 * *σ*_*G*_ + *m*. Assuming the parameters of the null model were correctly estimated, this results in a set false-positive rate of 0.13%, equivalent to a total of 195 cells falsely identified as gut-responsive out of a total of 150,000 cell segments identified in an experiment.

### Signal and noise correlation analysis

In order to compute noise and signal correlations between neuron in response to gut stimulation, we first extracted trial responses *x*_*it*_ for each gut-sensitive neuron *i* and trial number *t*. Signal correlations, 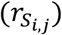, were then computed for each pair of neurons computing the pearson correlation between trial averaged responses: 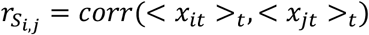 for neurons *i* and *j*, where <⋅>_*t*_ is the average over trials. Noise correlations between pairs were computed by averaging the correlation of each neuron’s trial residuals (trial response – averaged response) over all *T* trials: 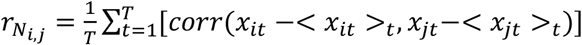. Matrices (as in **Fig. 4B,C**) were then constructed from these analyses by averaging all data from distinct neuron pairs within and between each brain region.

### Quantification and statistical analysis

Error bars displayed in the figures represent the standard error of the mean (SEM). Statistical analyses were performed using the SciPy Stats package.

