## Supplemental Information for "Whole-brain, all-optical interrogation of neuronal dynamics underlying gut interoception in zebrafish"

### Extended Data Figures

Extended Data 1

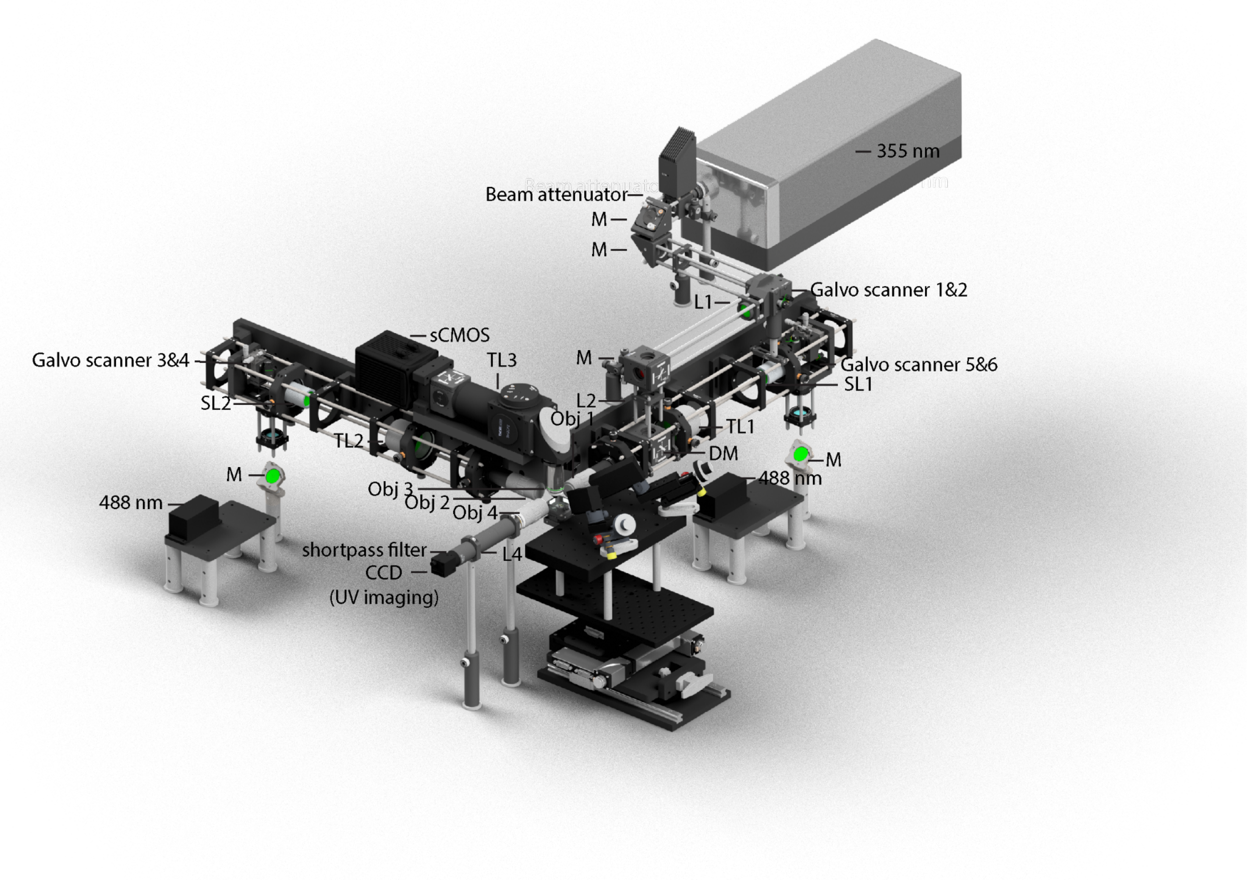


**Extended Data Fig. 1. Microscope setup.**

Schematic of the integrated UV uncaging path and light-sheet imaging system. A 3 W, 355 nm DPSS laser is attenuated by a custom beam attenuator at the laser head, then directed onto a pair of galvanometer scanners (1 & 2) for independent horizontal and vertical scanning. After passing through a telescope (L1 & L2), the UV beam is focused onto the sample via the light-sheet objective. A calibration pathway on the opposite side uses a lens (L4) and calibration camera to monitor the beam in real time, ensuring precise alignment and delivery of UV light to the gut. The light-sheet microscope is designed as described previously[17] and features a vertically mounted water-dipping detection objective (obj3, 16×/0.8 NA, Nikon) on a piezo stage, which projects fluorescence onto an sCMOS camera (Orca Flash 4.0, Hamamatsu). GCaMP emission is isolated by a 525/50 nm band-pass filter (525/50 nm, Semrock). Two horizontal illumination arms, each with an air objective (Obj 1 & 2, 4×/0.28 NA, Olympus), tube lens (TL 1 & 2), f-theta lens (SL 1 & 2), and galvanometer scanners (3-6, Cambridge Technology), generate the 488 nm excitation light-sheet, which is electronically shuttered to avoid stimulating the zebrafish eye. M, mirror; SL, scan lens; TL tube lens; DM, dichroic mirror; L, lens; Obj, objective.

Extended Data 2

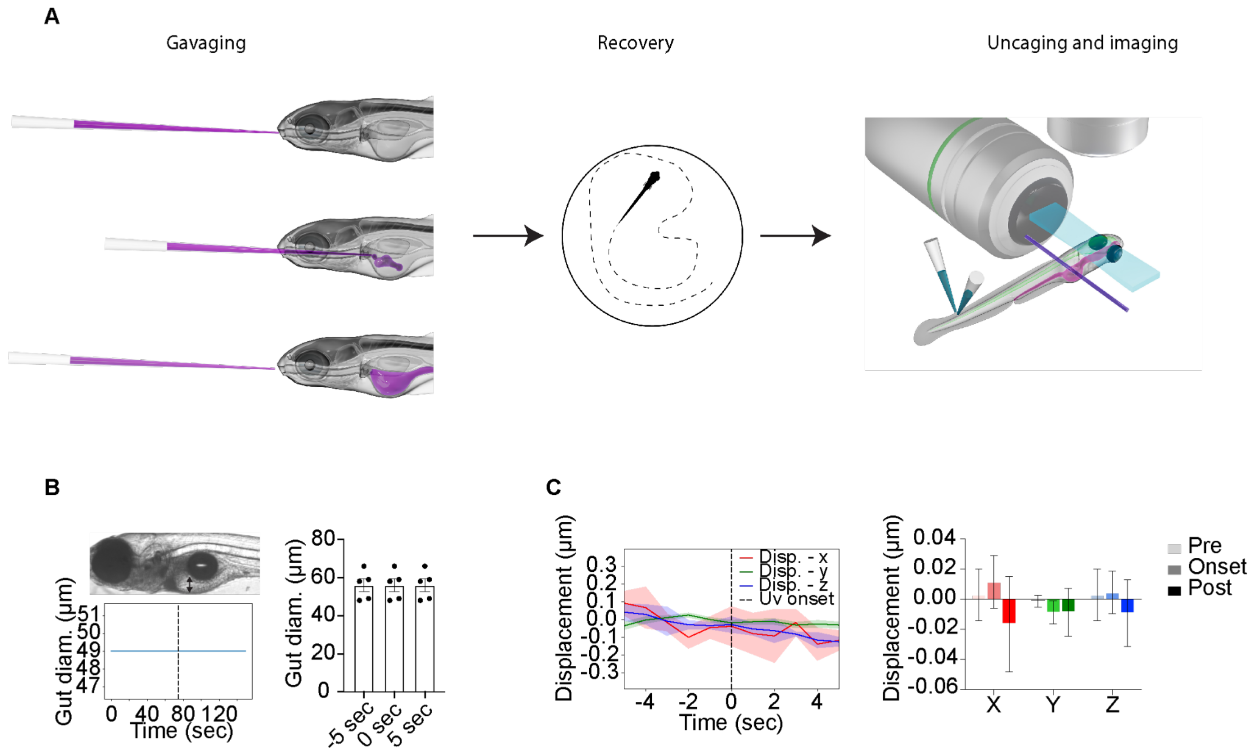


**Extended Data Fig. 2. Experimental overview.**

**(A)** Overview of the experimental procedure. After gavaging caged compounds into the larval zebrafish gut (left), the fish recover in a Petri dish (middle) for 5–10 minutes until they resume normal swimming behavior, before being transferred to the light-sheet microscope for uncaging and imaging (right).

**(B)** Chemical stimulation of the gut without mechanical dilation. Time-lapse measurements of gut diameter (left) show no discernible changes during uncaging, indicating minimal mechanical influence. Summary data (right) showing the gut diameter remains consistent before and after stimulation (mean ± SEM; N = 5 fish).

**(C)** Optical uncaging maintains tissue stability during imaging. Displacement in the x, y, and z axes (left) remains near baseline across pre-, onset-, and post-uncaging intervals, as shown by the summarized bar chart (right) (mean ± SEM; N = 5 fish).

Extended Data 3

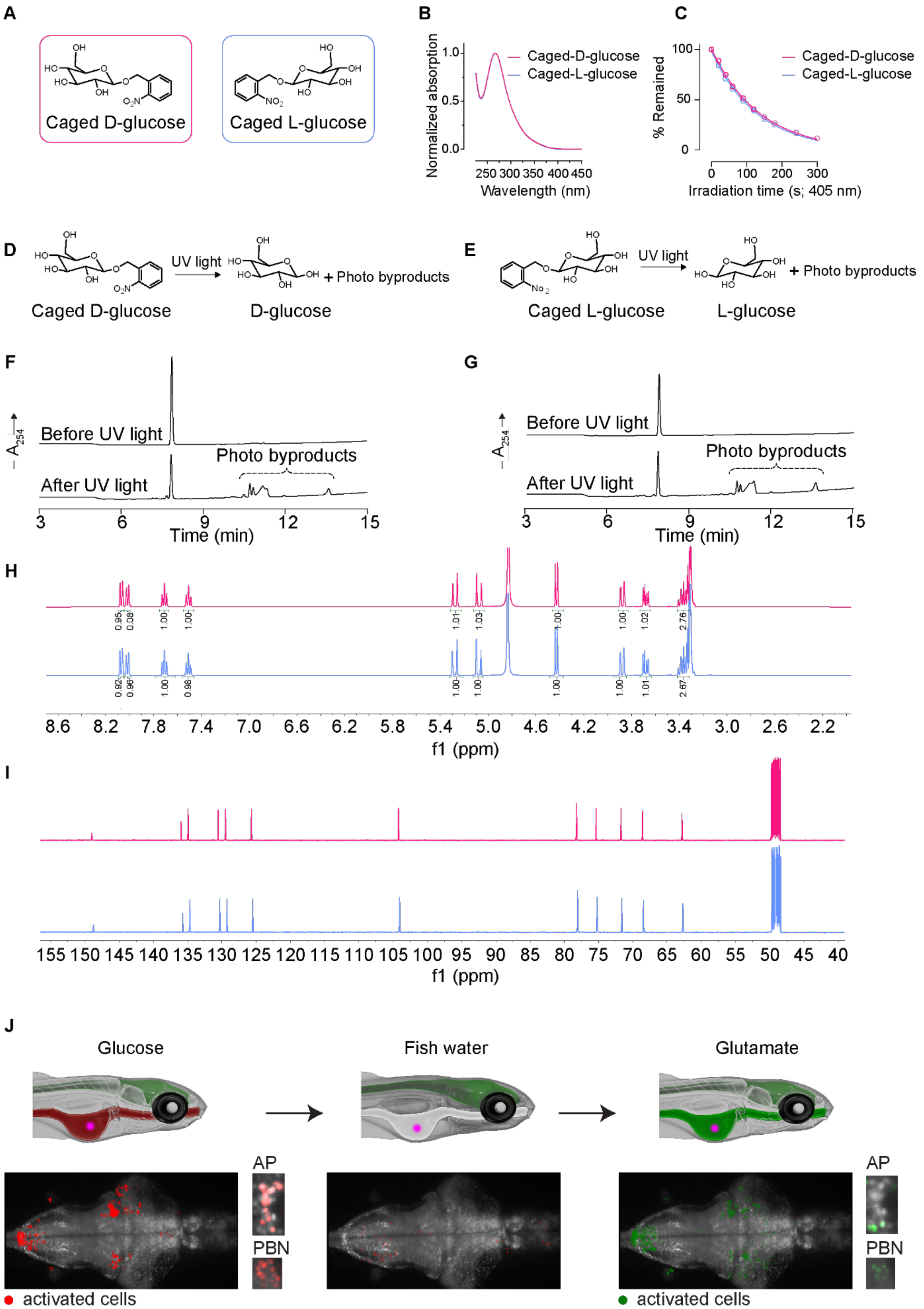


**Extended Data Fig. 3. Chemical structure and properties of caged D- and L-glucose.**

**(A)** Chemical structure of caged D-glucose and caged-L-glucose.
**(B)** Normalized absorption spectra for caged D- and L-glucose. Note that caged D- and L-glucose have effectively identical absorption spectra.
**(C)** Photolysis conversion vs. irradiation time for caged D- and L-glucose. Note that caged D- and L-glucose have effectively identical photolysis traces.
**(D)** Chemical uncaging operation for caged D-glucose
**(E)** Chemical uncaging operation for caged L-glucose.
**(F,G)** HPLC chromatograms of caged D- and L-glucose before and after irradiation. For experiments B-E, samples were prepared in PBS and irradiated with 405 nm LED (LOCTITE CL20 flood array). Chromatograms show caged-D-glucose and caged-glucose release of same photo byproducts.
**(H)** $^{1}$H NMR of caged-D-glucose (magenta) and caged-L-glucose (blue) recorded in $CD_3OD$.
**(I)** $^{13}$C NMR of caged-D-glucose (magenta) and caged-L-glucose (blue) recorded in $CD_3OD$. Note that caged-D-glucose and caged-L-glucose exhibited effectively identical $^{1}$H NMR spectra, $^{13}$C NMR spectra, indicating that the two chemicals share the same electronic environment. However, chirality measurement revealed a difference: caged-L-glucose had an optical rotation of $[\alpha]^{21}_{D}$ = 35±1° (c =0.1, pyridine), which is opposite of the literature value of caged-D-glucose, $[\alpha]^{21}_{D}$= −36.4° (c =0.55, pyridine). This opposite optical rotation confirms that the two chemicals are indeed an enantiomeric pair.
**(J)** Example case showing how multiple compounds can be introduced consecutively into the same fish to compare nutrients-evoked responses. In this example, glucose (left), fish water (middle), and glutamate (right) are sequentially delivered to the gut by microgavaging multiple times in the same fish.

Extended Data 4

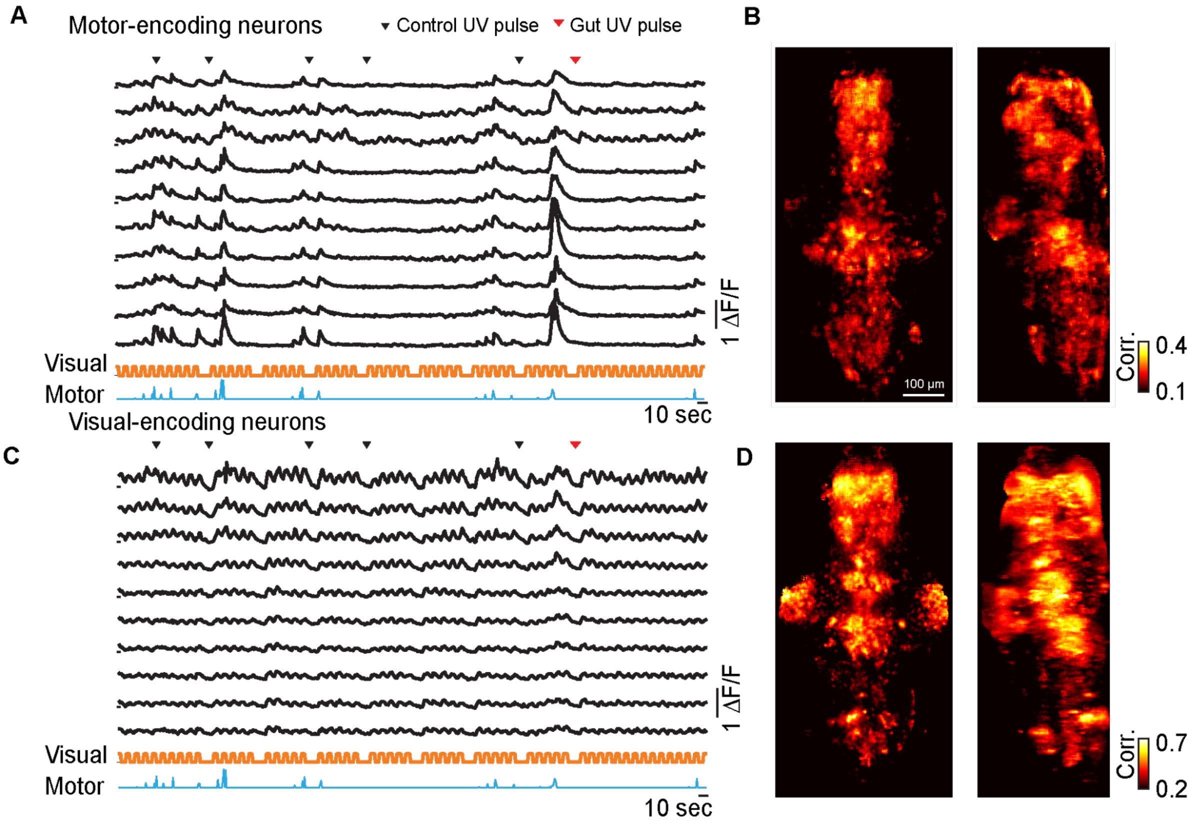


**Extended Data Fig. 4. Brain-wide detection of visual- and motor-responsive cells.**

**(A)** Example calcium traces of motor-responsive neurons. Each row represents the activity of a single neuron. The orange trace indicates visual motion, with upward segments representing forward movement of the visual gratings and downward segments indicating no motion. The blue trace reflects motor power derived from a sliding window of the tail voltage signal. The black triangles mark the onsets of control UV stimulation (UV targeted outside of the fish), and the red triangle indicates onset of gut stimulation.

**(B)** Average brain map showing the spatial location of motor-related neurons. Left: dorsal view. Right: sagittal view. Colors represent the correlation coefficient between neuronal activity and motor signals (N = 6 fish).

**(C)** Example calcium traces of visual-related neurons.

**(D)** Average brain map showing the spatial location of visual-responsive neurons (N = 6 fish).

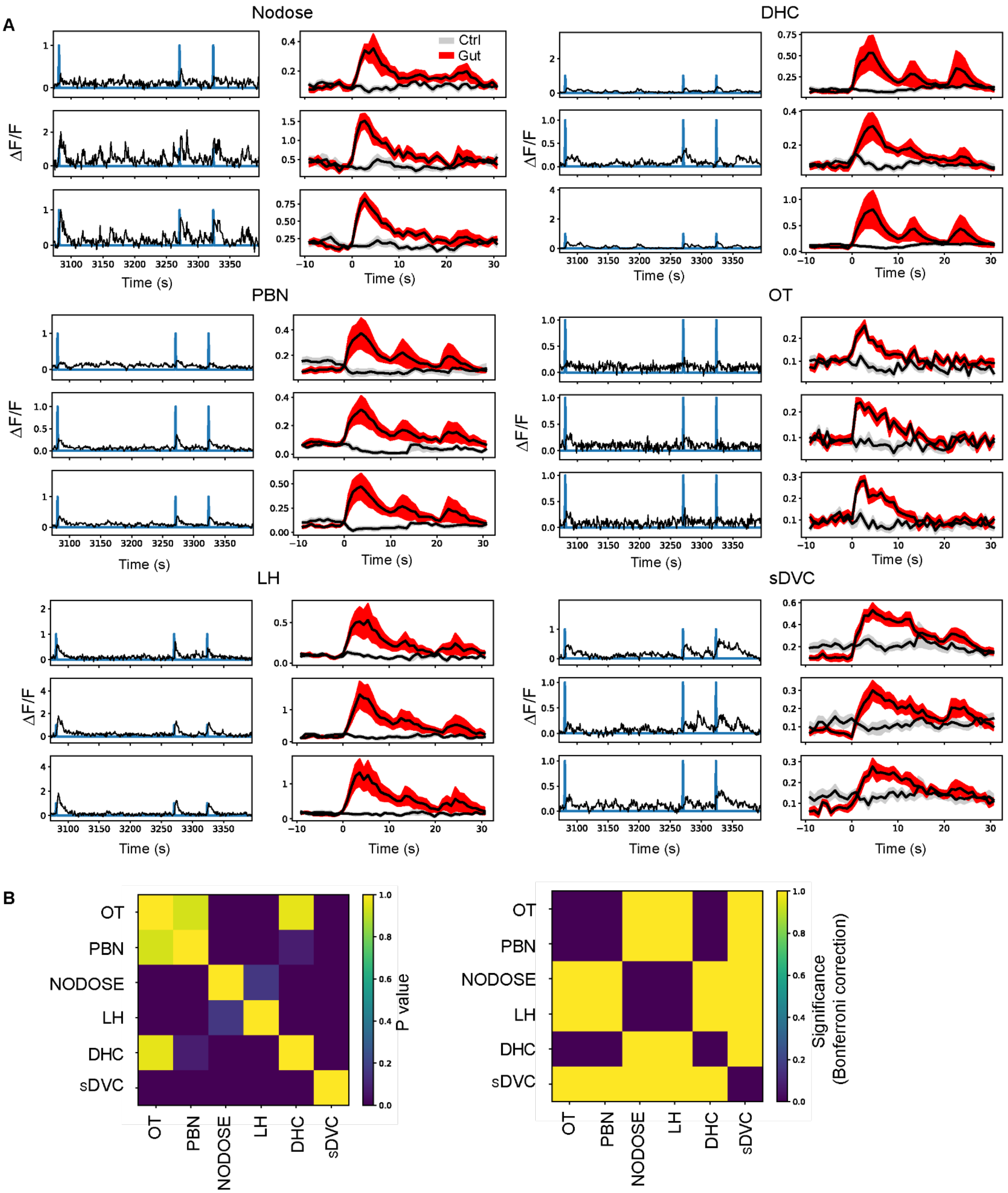


**Extended Data Fig. 5. Noise Correlation Traces and Timescale Significances.**
**(A)** Example time series (left) and averages (right) to gut and control UV pulses for a selection of cells in various brain regions. Note that nodose, more than any other brain region, contains neurons with gut responses consisting of multiple events. Pooling from multiple units may smooth these responses in the brain, reducing noise correlations between nodose cells and cells within the brain, as seen in Fig. 4C.

**(B)** Matrix of p-values for rank sum tests of differences in FWHM, as seen in Fig. 4F (left), and results of Mann-Whitney significance tests after Bonferonni correction (right). Values of elements equal to one indicate statistical significance.

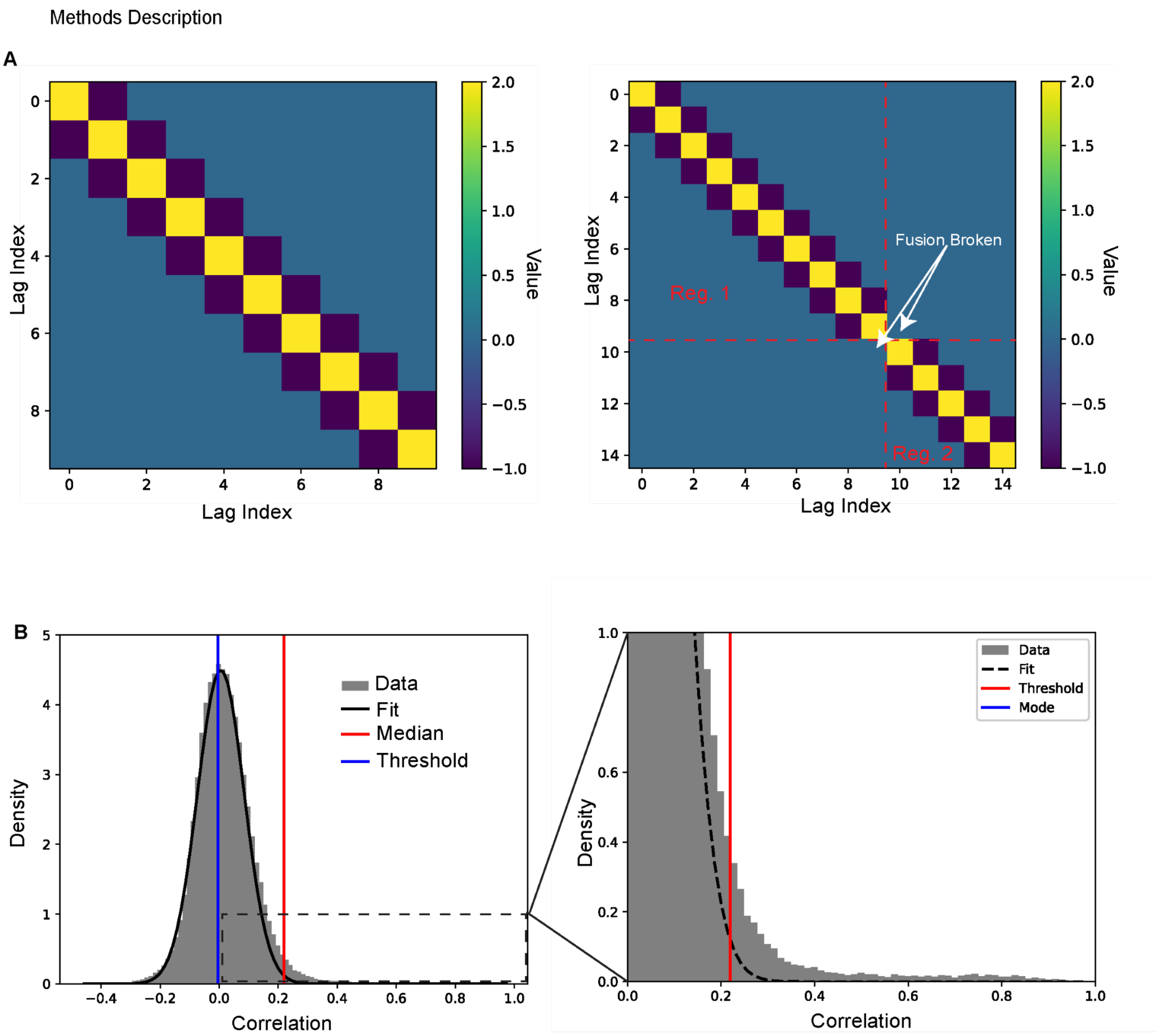


**Extended Data Fig 6. Methods demonstration.**

**(A)** Left: a finite difference matrix for a single regressor of lag order 10. Right: a modified finite difference matrix for two regressors (shown as boxes in red dashed lines). Regressor 1 has a lag order of 10, and regressor 2 has a lag order of 5. Fusion between the two regressors is broken by setting $D_{b_j,b_j+1} = 0$ and $D_{b_j+1,b_j} = 0$ for all boundary indices $b_j$.

**(B)** Left: the distribution of correlations between each neuron and its best fit linear model (gray) was used to construct a null model for assessing whether a neuron has a statistically significant response to gut stimulation. Data from the left of the mode (blue line) was used to fit the left side of a Gaussian distribution (black line). The threshold for significant responses was then set to the mode plus three times the estimated standard deviation (red line). Right: blow-up of the box in left highlighting the presence of neuron correlations above the computed threshold.
